# A semi-parametric Bayesian approach, iSBA, for differential expression analysis of RNA-seq data

**DOI:** 10.1101/558270

**Authors:** Ran Bi, Peng Liu

**Affiliations:** Department of Statistics, Iowa State University, Ames, Iowa, USA

## Abstract

RNA sequencing (RNA-seq) technologies have been popularly applied to study gene expression in recent years. Identifying differentially expressed (DE) genes across treatments is one of the major steps in RNA-seq data analysis. Most differential expression analysis methods rely on parametric assumptions, and it is not guaranteed that these assumptions are appropriate for real data analysis. In this paper, we develop a semi-parametric Bayesian approach for differential expression analysis. More specifically, we model the RNA-seq count data with a Poisson-Gamma mixture model, and propose a Bayesian mixture modeling procedure with a Dirichlet process as the prior model for the distribution of fold changes between the two treatment means. We develop Markov chain Monte Carlo (MCMC) posterior simulation using Metropolis Hastings algorithm to generate posterior samples for differential expression analysis while controlling false discovery rate. Simulation results demonstrate that our proposed method outperforms other popular methods used for detecting DE genes.

## Introduction

During the past decade, RNA sequencing (RNA-seq) technologies have revolutionized transcriptomic studies. In a typical RNA-seq experiment, messenger RNA (mRNA) molecules are extracted from samples, fragmented, and converted to a library of complementary DNA (cDNA) fragments. The cDNA fragments are then amplified and sequenced on a high-throughput platform, such as HiSeq by Illumina or SOLiD by Applied Biosystems. Millions of DNA fragment sequences, called reads, are obtained for each sample and mapped to a reference genome. The number of reads aligned to a given gene measures the expression level for that gene. Thus, RNA-seq generates discrete count data rather than continuous data serving as measurements of mRNA expression levels.

In the statistical analysis of RNA-seq data, detecting differentially expressed (DE) genes across treatments or conditions is one of the major steps and often the main goal. A gene is considered to be DE if the expression levels change across treatment groups. Otherwise, the gene is said to be equivalently expressed (EE). Generally, negative binomial (NB) distribution is used for modeling RNA-seq count data. Many statistical methods based on the NB distribution have been proposed for detecting DE genes with RNA-seq data, including *edgeR* [1–4], DESeq [5] and *DESeq2* [6]. Methods that do not assume NB models typically involves transformation of the count data to continuous scale, such as the *Voom* and *limma* pipeline [7], which models the mean-variance relationship of the log-transformed count data and produces a precision weight for each observation, then applies the *limma* method based on normal distributions [8] for the detection of DE genes.

The comparison among all the popular methods for RNA-seq data analysis mentioned above has been done through simulation studies [9, 10]. However, the optimality of these existing testing procedures is inadequately studied. Si and Liu (2013) [11] developed an optimal test for RNA-seq data anaysis while controlling FDR, where optimal tests were defined as tests that achieve the maximum of the power averaged across all genes for which null hypotheses are false. Furthermore, Si and Liu (2013) [11] proposed an approximation to the optimal test, where hyper distributions were estimated with mixture distributions, and such a test is called the approximated most average powerful (AMAP) test. In the two-treatment comparison problem, Si and Liu (2013) [11] modeled the gene-specific treatment means by the overall geometric mean expression level across both treatments and the ratio of the two treatment means, i.e., fold change *ρ*_*g*_. They used a *K*-component mixture Gamma-Normal (MGN) distribution to model the joint distribution of the overall geometric mean expression level and the logarithm of the fold change. However, there are several limitations of using MGN distribution, such as difficulty in selecting an appropriate number of components *K*, and challenges in modeling the empirical distribution of all genes by parametric models.

Bayesian nonparametric modeling is a more flexible way for distribution estimation and is often applied to avoid critical dependence on parametric assumptions. The most popular Bayesian nonparametric methods adopt Dirichlet process (DP) mixture modeling, and such modeling framework has been utilized for DE analyses. For instance, [12] chose DP mixtures to model the population of genes under two different conditions and applied to a microarray dataset. Liu *et al.* (2015) [13] used the DP prior for modeling the distribution of fold changes between two treatments, with a mixture of a point mass at one and a Gamma distribution as the base distribution in the DP prior. In the method proposed by Liu *et al.* (2015) [13], one treatment condition was set as the reference condition (i.e., baseline) and they used DP as the prior for the distribution of fold changes of the other condition versus the reference. When they changed the reference treatment group, the declared differential expression status were not exactly the same for all genes.

To address this issue that the model is not invariant to the choice of reference condition, we propose a method using a mixture of three components as the base distribution in the DP prior for the distribution of the fold changes between two treatment conditions. The three components are a point mass at one, a Gamma, and an inverse-Gamma distribution, so that the model becomes invariant no matter which treatment group is set to be the reference. In addition, we model RNA-seq count data via a Poisson-Gamma mixture model, which is equivalent to a NB model. Similar to Liu *et al.* (2015) [13], this paper shows how our mixture modeling procedure can be accommodated to provide meaningful posterior probabilities of simple or composite null hypothesis. Also, we show that the posterior inference can be viewed as an approximation for the optimal test in Si and Liu (2013) [11], thus our approach is an approximated optimal test.

The article is organized as follows. In the Methods section, we describe our proposed Bayesian mixture modeling pipeline and the prior models, then present the MCMC sampling scheme for posterior inference and FDR estimation. In the Results section, we generate several simulation studies based on NB distributions, and compare our proposed method to some popular methods for DE analysis. We also analyze a real dataset using our proposed method. The Discussion section summarizes our results and provides some discussion.

## Methods

In this section, we first describe the framework of our mixture modeling, and then introduce the prior models employed in our method.

### A Poisson-Gamma Mixture Model

Suppose that an RNA-seq experiment measures *G* genes. Let *Y*_*gij*_ denote the number of reads mapped to gene *g* from biological replicate *j* of treatment *i*, where *g* = 1, *…, G, i* = 1, 2, *j* = 1, *…, n*_*i*_, and *n*_*i*_ is the number of biological replicates in treatment *i*. As we mentioned in the introduction section, NB distribution has been popularly applied to such data. In the development of our modeling framework, we use a Poisson-Gamma mixture model parameterization instead of the NB model directly, where the RNA-seq read counts follow a Poisson distribution conditioning on the true expression mean, and the true gene abundances follow a Gamma distribution between replicate RNA samples. Then read count data *Y*_*gij*_ can be modeled as below,

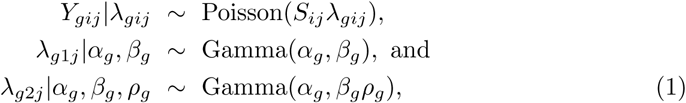

where *S*_*ij*_ is a normalization factor that accounts for sequencing depth variation and nuisance technical effects across the replicates, *λ*_*gij*_ is the normalized expression mean of *j*th replicate of *i*th treatment in gene *g, α*_*g*_ is the shape parameter which stands for the reciprocal of the dispersion parameter for gene *g, β*_*g*_ is the rate parameter for the first treatment, and the product of *β*_*g*_ and *ρ*_*g*_ is the rate parameter for the second treatment. So the marginal expression mean for treatment 1 is *α*_*g*_*/β*_*g*_, while for treatment 2 is *α*_*g*_*/*(*β*_*g*_*ρ*_*gi*_). Therefore, the mean ratio of treatment 1 over treatment 2 is *ρ*_*g*_, which refers to the fold change between treatment 1 versus treatment 2.

The goal of differential expression analysis is to test

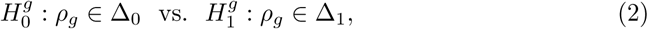

for each gene *g*, where Δ_0_ represents the null set of values for *ρ*_*g*_, while Δ_1_ represents the alternative set. Δ_0_ and Δ_1_ are assumed to be a partition of the positive real line ℝ^+^ (Δ_0_ ⋃ Δ_1_ = ℝ^+^, Δ_0_ ⋂ Δ_1_=Ø). The null space Δ_0_ can be defined in different ways depending on the biological problems of interest. For example, if we are interested in identifying DE genes across the two treatments, we set Δ_0_ = {1}. If we are interested in whether the mean expression level in the first treatment is greater than the second treatment, we set Δ_0_ = (0, 1]. If we are interested in genes whose expression changes are large enough, for instance, the fold changes are greater than 1.5 [14], we set Δ_0_ = [1*/*1.5, 1.5].

### Prior Specification

Since our main focus is to test the hypothesis about the fold change parameter *ρ*_*g*_ in (2) for each gene, specifying an appropriate prior distribution for *ρ*_*g*_ is very crucial. The empirical distribution of the fold change of all genes could be very irregular and differs between various studies. To provide maximal flexibility, Bayesian nonparametric modeling with DP is a common way for distribution estimation. DP is a stochastic process whose realizations are probability distributions, i.e., each draw from a DP is itself a distribution. The formal definition of DP is as follows. Given a measurable set Ω, a base probability distribution *F*_0_ and a positive real number *M* called the concentration parameter, a random probability distribution *F* is generated by a DP if for any measurable partition *A*_1_, *…, A*_*k*_ of Ω, the distribution of (*F* (*A*_1_), *…, F* (*A*_*k*_)) is Dirichlet *D*(*M · F*_0_(*A*_1_), *…, M · F*_0_(*A*_*k*_)). We denote this by *F ∼ DP* (*M, F*_0_). The parameters *F*_0_ and *M* play intuitive roles in the definition of the DP. For any measurable subset *B* of Ω, the base distribution *F*_0_ is the mean of the DP, i.e., *E*[*F* (*B*)] = *F*_0_(*B*). Besides, the concentration parameter *M* defines the variance as *V ar*[*F* (*B*)] = *F*_0_(*B*)(1*-F*_0_(*B*))*/*(*M* + 1). The larger *M* is, the smaller the variance, and the DP will concentrate more of its mass around the mean.

Throughout our mixture modeling procedure, we use a DP to model the fold change parameters (*ρ*_1_, *…, ρ*_*G*_). Different from Liu *et al.* (2015) [13], we use a mixture of a point mass at one, a Gamma and an inverse-Gamma distribution as the base distribution in the DP prior for the distribution of the fold change parameters, so that our modeling is invariant to the specification of the reference condition, and we call it *iSBA* (where *SBA* stands for semiparametric Bayesian approach). Details of the proof of reference level invariance are provided in S1 Appendix.

Therefore, the DP prior for gene *g, g* = 1, *…, G*, can be expressed as

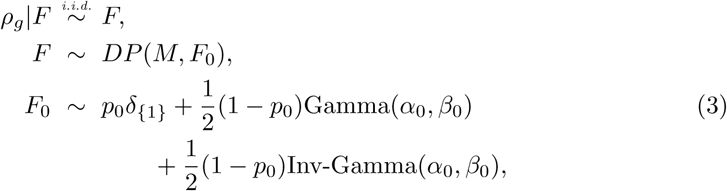

where *p*_0_ is the proportion of EE genes, and *δ*_*{x}*_ denotes a point mass at *x*. In this paper, we set *p*_0_ = 0.5 to give no prior preference to either DE or EE. The concentration parameter *M* in the DP priors is fixed as *M* = 1, which is a common choice used in application [12, 15, 16]. Throughout our paper, the simple null hypothesis of our great interest is *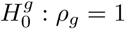*.

Following Liu *et al.* (2015) [13], we use a Gamma distribution as the prior distribution for *β*_*g*_ due to its conjugacy, and an exponential distribution as the prior for *α*_*g*_ in order to reduce the computational complexity of the posterior distribution,

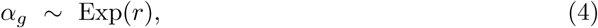

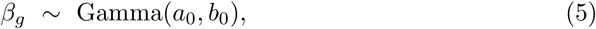

where *r, a*_0_, *b*_0_, in addition to *α*_0_ and *β*_0_ in (3), are hyperparameters. We set *r* = 0.01, *a*_0_ = 0.1, *b*_0_ = 0.1, *α*_0_ = 0.1, *β*_0_ = 0.1 so that the priors are non-informative and the inference for *α*_*g*_ and *β*_*g*_ mainly relies on the observed data. For computational simplicity, we set the priors for *α*_*g*_’s, *β*_*g*_’s, and *ρ*_*g*_’s to be independent.

### Markov Chain Monte Carlo Simulation

Posterior inference based on our proposed model is implemented by using Markov chain Monte Carlo (MCMC) algorithm [17]. MCMC methods are usually employed to generate samples from the posterior distribution by constructing a Markov chain that has the target posterior distribution as its equilibrium distribution. We use an MCMC-based sampling method in our proposed Bayesian mixture models. Gibbs sampling is the most frequently used tool to perform MCMC algorithm for Bayesian hierarchical models when dealing with conjugate priors. However, for addressing non-conjugate priors, the simplest way is by using the Metropolis-Hastings algorithm [18].

The Metropolis-Hastings algorithm simulates samples from a target distribution *π*(*x*) using a proposal distribution *g*(*x\**|*x*), and updates the state *x* as follows. Generate a candidate state *x\** from the distribution *g*(*x\**|*x*), then compute the acceptance probability

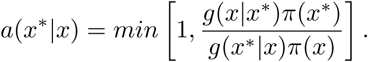

Set the new state *x′* to *x\** with probability *a*(*x* x*). Otherwise, reject the candidate *x\**and let *x′* be the same as *x*.

To simplify the use of DP prior, when *F* is integrated over its prior distribution (3), the sequence of *ρ*_*g*_’s follows a Polya urn scheme [19, 20], that is,

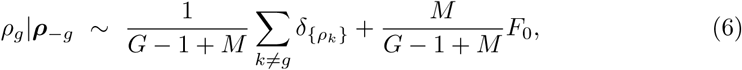

where ***ρ***_*-g*_ is the vector of (*ρ*_1_, *…, ρ*_*G*_) after deleting *ρ*_*g*_.

Then the most direct approach to sample for our model is to perform Metropolis-Hastings update for each of the *ρ*_*g*_. However, this algorithm may not be very efficient since it cannot change the *ρ*_*g*_ for more than one gene simultaneously. A change to the *ρ*_*g*_ values occurs only when they are reallocated to new components. Thus it may take long time to converge to the posterior distribution [21]. In order to improve the efficiency of the MCMC algorithm, a modified Metropolis-Hastings updates and partial Gibbs sampling method has been proposed by Neal (2000) [21] (Algorithm 7). Suppose *K* is the number of distinct values in the vector (*ρ*_1_, *…, ρ*_*G*_) and the distinct values are denoted as *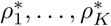*, respectively. Let ***ξ*** = (*ξ*_1_, *…, ξ*_*G*_) be the configuration indicators defined by

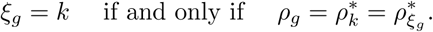

Therefore, we reparameterize the prior model for *ρ*_*g*_’s with *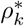*’s and *ξ*_*g*_’s as follows,

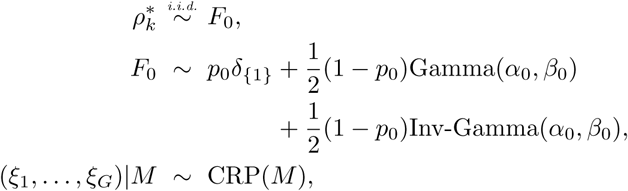

where the prior models for *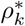* ‘s and *ξ*_*g*_’s are independent and CRP stands for Chinese Restaurant Process. CRP is a random distribution and the full conditional distribution for *ξ*_*g*_’s can be written as

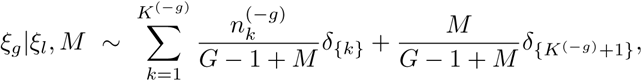

where *K*^(*-g*)^ denotes the number of distinct values in the vector (*ρ*_1_, *…, ρ*_*G*_) after deleting *ρ*_*g*_, and 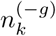 denotes the number of (*ρ*_1_, *…, ρ*_*G*_) who equal *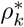* after deleting *ρ*_*g*_.

The MCMC sampling scheme uses the modified Metropolis-Hastings updates and partial Gibbs sampling method to repeatedly sample the following parameters step by step. The procedure for generating the full conditionals of all parameters and how we apply Metropolis-Hastings algorithm are shown in S2 Appendix.

1. Draw samples of *λ*_*gij*_’s from their full condition distributions,

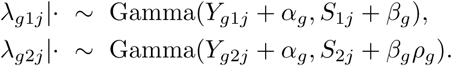
2. Draw samples of *β*_*g*_’s from their full conditional distributions,

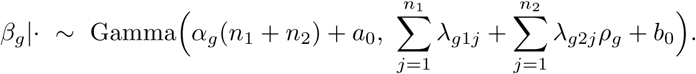
3. There is no closed-form full conditional distribution for *α*_*g*_’s. Since the conditional posterior distribution for each gene *g* is a log-concave function with respect to *α*_*g*_, we could draw posterior samples based on adaptive rejection sampling method [22].
4. Obtain posterior samples for *ρ*_*g*_’s by getting the Markov chain for (*ξ*_1_, *…, ξ*_*G*_) and 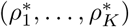 as follows:
  i. Update the configuration vector (*ξ*_1_, *…, ξ*_*G*_).
    - For *g* = 1, *…, G*, repeat the following: If *ξ*_*g*_ = *ξ*_*l*_ for some *l ≠ g*, let 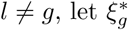 be a newly created component, with 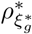 drawn from *F*_0_. Set *ξ*_*g*_ to 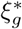 with probability

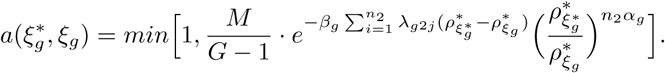

Otherwise, if *ξ*_*g*_ *≠ ξ*_*l*_ for all *l ≠ g*, draw 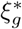 from ***ξ***_*-g*_, choosing 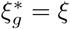 with probability 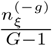. Set the new *ξ*_*g*_ to this 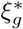 with probability

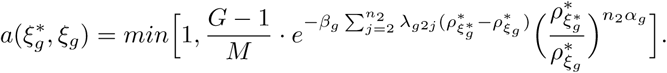
    - For *g* = 1, *…, G*, if *ξ*_*g*_ *≠ ξ*_*l*_ for all *l ≠ g*, do nothing. Otherwise, choose a new value for *ξ*_*g*_ from {*ξ*_1_, *…, ξ*_*G*_} with probabilities

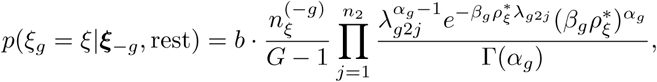

where *b* is the appropriate normalizing constant.
  ii. Update 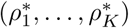. For *k* = 1, *…, K*, repeat the following: Draw 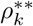 from *F*_0_. Set the new value of 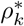 to 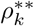 with the probability

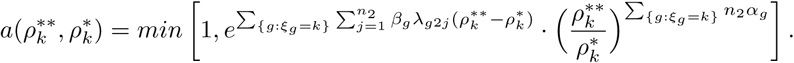 Otherwise, let the new 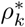 be the same as the old value. If we have duplicated 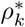, delete it and combine *ξ*_*g*_.

### Bayesian FDR Control

In genomic studies, tens of thousands of hypotheses are simultaneously tested, each relating to a gene. Thus multiple testing procedures that control the number of false significant results are commonly used in the analysis. False discovery rate (FDR) [23], defined as the expected proportion of false positives among the rejected hypotheses, has been the common choice of error criterion in RNA-seq data analysis. Within the Bayesian framework, one can estimate the FDR with Bayesian FDR [24, 25] by using posterior probability.

For each gene *g, g* = 1, *…, G*, the posterior probability that *g*th null hypothesis is true is denoted by *P* (*ρ*_*g*_ *∈* Δ_0_*|****Y***_*g*_). If we are interested in detecting DE genes, with Δ_0_ = {1}, *P* (*ρ*_*g*_ *∈* Δ_0_*|****Y***_*g*_) is the posterior probability that gene *g* is EE. *P* (*ρ*_*g*_ *∈* Δ_0_*|****Y***_*g*_) can be estimated by the proportion of the posterior samples obtained from MCMC for gene *g* that fall into the null set Δ_0_, i.e.,

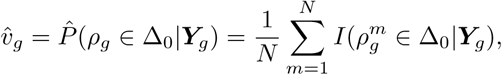

where *N* is the number of posterior samples. We reject 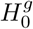 if the estimated posterior probability 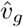 is smaller than a critical value *c\**. The critical value *c\** is chosen based on controlling the FDR at a target level *γ*, for example, 0.05, i.e.,

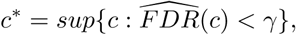

where

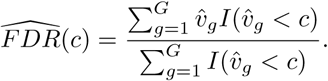

So the Bayesian FDR controlled at level *γ* can be calculated by

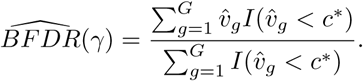

## Results

In this section, we adopt the simulation settings in Liu *et al.* (2015) [13] to assess the performance of our proposed method (*iSBA*), and compare to their semi-parametric Bayesian (*SBA*) method along with other popular methods for differential expression analysis of RNA-seq data, such as *edgeR* [3], *voom* and *limma* pipeline [7], and *DESeq* [5]. To mimic the distributions of real RNA-seq count data, we sampled gene-specific mean and dispersion parameters from the estimated values based on a maize study [26] that compared gene expression between bundle sheath and mesophyll cells of corn plants. Following Liu *et al.* (2015) [13], we conducted the same two sets of simulation studies (A and B). For each simulation study, 32 independent RNA-seq datasets were simulated from NB distributions with given mean and dispersion parameters, each dataset contains 10,000 genes, 2 treatment groups, and *n* replicates per treatment group, where *n* = 3 or 6. For our proposed method, we generated 5000 posterior samples after 3000 iterations burn-in, to calculate the estimated posterior probabilities. Convergence was checked via Gelman-Rubin criteria [27]. The test performances of different methods are evaluated by averaging the 32 datasets.

### Simulation A

We used the maize dataset published by Tausta *et al.* (2014) [26] to estimate the gene-specific mean for one treatment group and the dispersion parameters, and randomly sampled 10,000 pairs of mean and dispersion parameters out of all 27,819 pairs without replacement, which would be used as the true mean expression level for the control group (*µ*_*g*_) and the true dispersion parameter (*ϕ*_*g*_) for gene *g* = 1, *…*, 10000. Given the number of replicates per treatment group, *n* = 3 or 6, the RNA-seq read count data for the control group were generated from *NB*(*µ*_*g*_, *ϕ*_*g*_) for gene *g*. Then we randomly selected 5000 out of the 10,000 genes to be EE, whose count data for the treatment group were also drawn from *NB*(*µ*_*g*_, *ϕ*_*g*_). The remaining 5000 genes were simulated to be DE genes, with fold change (*ρ*_*g*_) set to be 4, 8, 0.25 and 0.125. Thus we had 1250 genes for each *ρ*_*g*_ value, whose count data for the treatment group were drawn from *NB*(*µ*_*g*_*ρ*_*g*_, *ϕ*_*g*_).

### Simulation B

Similar to Simulation A, we generated 10,000 genes from *NB*(*µ*_*g*_, *ϕ*_*g*_), with fold change *ρ*_*g*_ for 5000 DE genes. Instead of setting *ρ*_*g*_ to be 4, 8, 0.25 or 0.125, we simulated *ρ*_*g*_ from a two-component mixture of lognormal distributions,

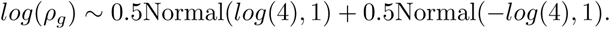

### Simulation Results for Testing DE Genes

In order to avoid the impact on test performance with different normalization procedures, we applied the same normalization steps for all the methods under comparison. Specifically, we set all normalization factors to be 1 for both Simulations A and B.

The receiver operating characteristic (ROC) curves that plot the true positive rate (TPR) versus false positive rate (FPR) resulting from Simulations A and B with number of replicates per group *n* = 3 or 6 are shown in Fig 1. These curves were generated based on either the posterior probabilities or *p-*values for each method. For each level of FPR, the TPRs were averaged over the 32 simulated datasets. We plotted the curves over the FPR values in the range of 0 and 0.1 because we are most interested in small FPR values. We also calculated the area under the curve (AUC) values as the percentages of 0.1, which is the total area in the range of FPR *<* 0.1. The average values and standard deviations of the AUC across the 32 simulated datasets are reported in the legends of Fig 1. Fig 1 shows that our *iSBA* method and the *SBA* method proposed by Liu *et al.* (2015) [13] generated the highest ROC curves and largest AUC values among all tests under all simulation settings, indicating that *iSBA* and SBA methods outperformed other methods in terms of ranking DE genes.

**Fig 1.**
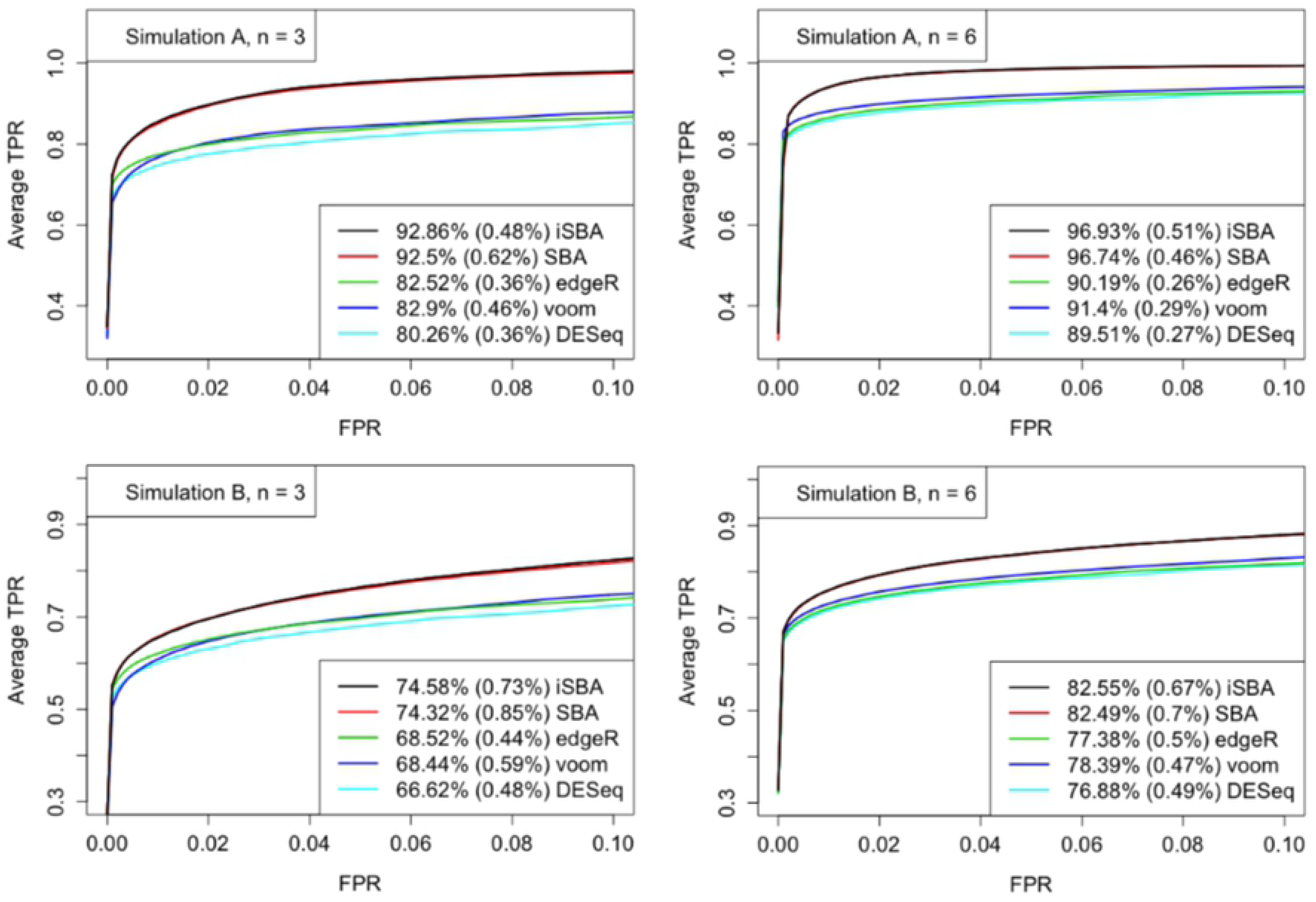
ROC curves resulting from Simulations A and B. For each level of FPR, the TPRs were averaged over the 32 simulated datasets. The percentage reported in the legend is the average AUC for each method, representing the percentage of 0.1, which is the total area in the range of FPR *<* 0.1, and the percentage in each set of parentheses is the standard deviation of the estimated AUC.

We also checked the false discovery (FD) plot as in Liu *et al.* (2015) [13], which is the plot of the number of false positives versus the number of top ranked genes selected as DE. Genes were ranked based on either posterior probabilities or *p-*values for each method. A better performing method would have a lower FD curve. The FD plots for Simulations A and B with *n* = 3 or 6 are shown in Fig 2. The number of false positives decreased when sample size increased from 3 to 6 for all methods, as expected. Our *iSBA* method and the *SBA* method provided the lowest FD curves under all simulation settings, indicating that our *iSBA* method and the *SBA* method produced less false positives than others, when we declared the same number of DE genes for all methods.

**Fig 2.**
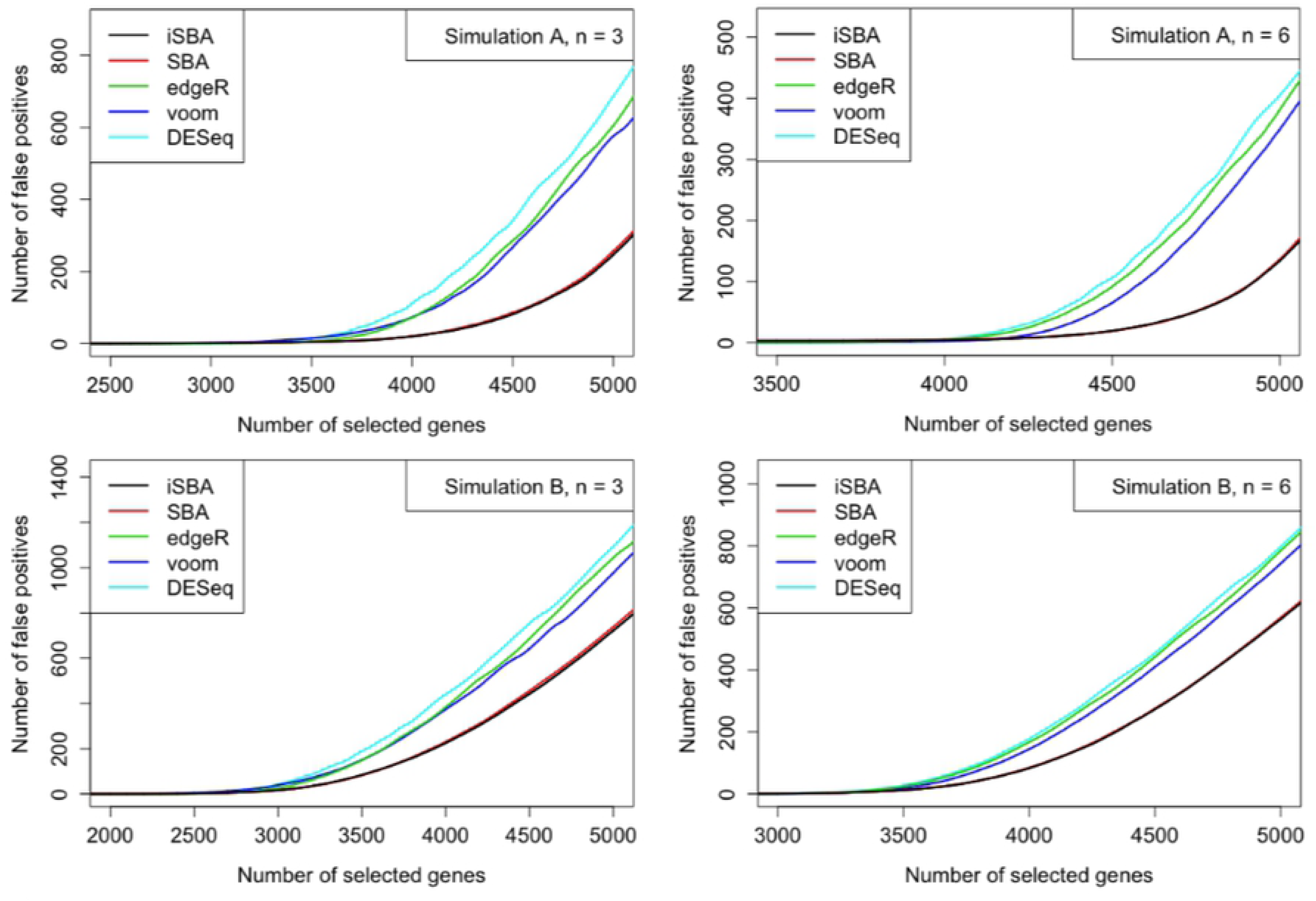
False discovery curves resulting from Simulations A and B. For each number of top ranked genes selected as DE, the number of false positives were averaged across the 32 simulated datasets. Genes were ranked based on either posterior probabilities or *p-*values.

In addition, we evaluated the estimation of FDR based on subsection “Bayesian FDR Control” in Methods Section for our method and *SBA* method. For other non-Bayesian methods, we applied the Benjamini and Hochberg [23] procedure to adjust *p*-values for multiple comparisons. FDR plots for Simulations A and B with *n* = 3 or 6are presented in Fig 3. Our *iSBA* method controlled FDR well, the *SBA* method performed the second, while other methods provide more conservative results.

**Fig 3.**
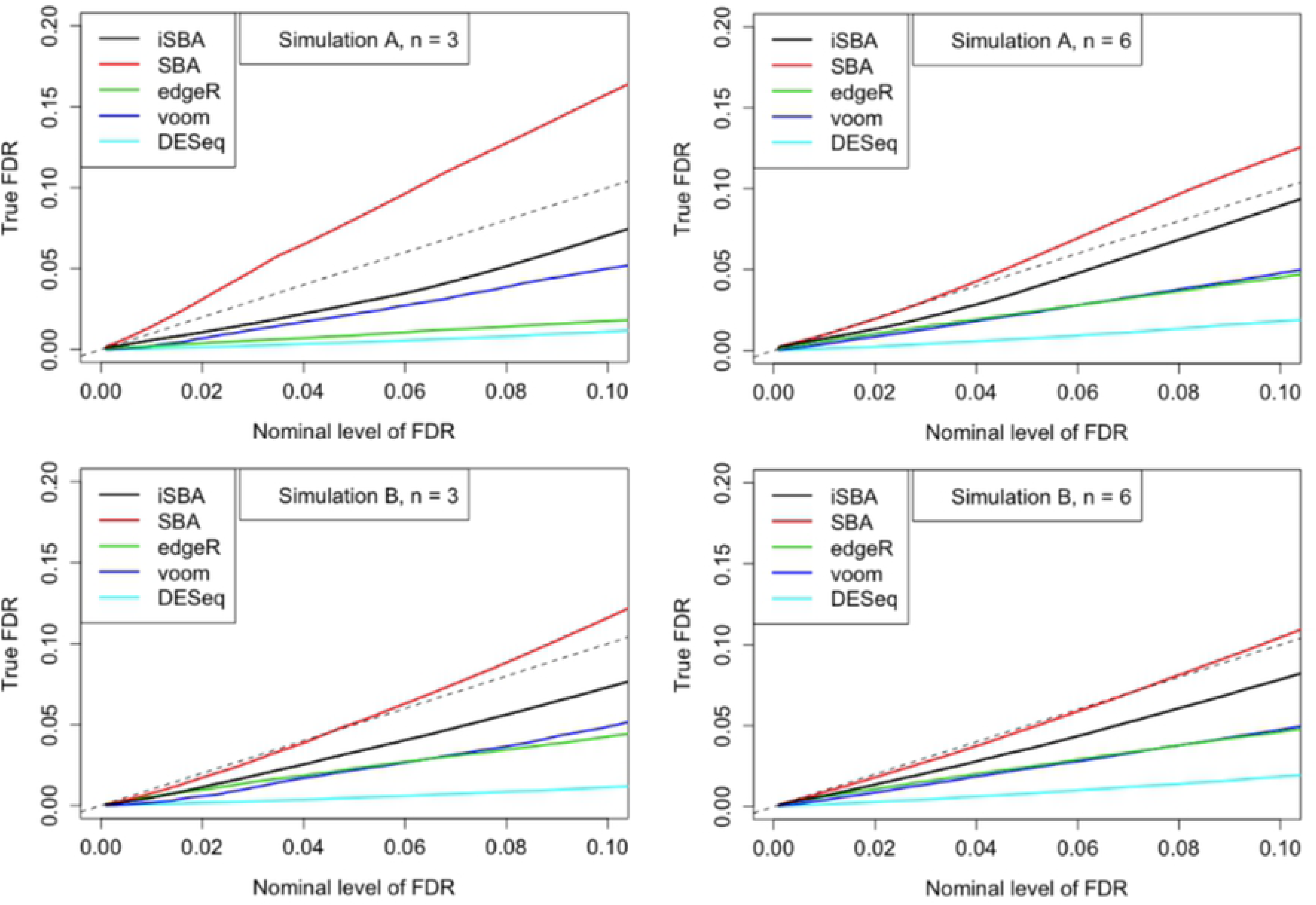
Plots of the actual FDR versus the nominal level of FDR resulting from Simulations A and B. The dashed lines correspond to the *Y* = *X* line. A well performing method would control the FDR below or close to the dashed line.

Based on results from these simulations, our *iSBA* method and the *SBA* method generated the highest ROC curves and the least false positives, comparing with other popularly applied RNA-seq DE analysis methods. Furthermore, the *iSBA* method controlled FDR the best, hence provided reliable lists of declared DE genes. All in all, our proposed *iSBA* method worked the best or among the best under all simulation settings.

### Simulation Results for Testing: *|logFC| ≤ log*1.5

In addition to the simple hypothesis testing problem introduced in the last subsection, we could also apply our method to do other types of hypothesis testing, for example, testing whether the fold change falls into a certain interval or not. In practice, biologists often want to detect genes whose fold-changes are big enough and biologically meaningful. This subsection shows the results for testing: *|logFC| ≤ log*1.5 for Simulation B.

We applied our *iSBA* method and the *SBA* method directly to do this hypothesis testing problem. For other methods including *edgeR, voom* and *limma* pipeline, and *DESeq*, we adopted the two-step procedure described in [11]. More specifically, in the first step, *ρ*_*g*_ = 1 was tested for each gene, and a list of DE genes was identified while controlling FDR at a given level. In the second step, among those DE genes declared in the first step, genes with large enough fold changes (*|logFC| > log*1.5) were selected.

The ROC curves for testing *|logFC| ≤ log*1.5 for Simulation B are shown in the upper panel of Fig 4. The *iSBA* method and the *SBA* method outperformed all other methods. The lower panel of Fig 4 provides the FDR plots, from which we could notice that our *iSBA* method controlled FDR well in the range of FDR smaller than 0.1.

**Fig 4.**
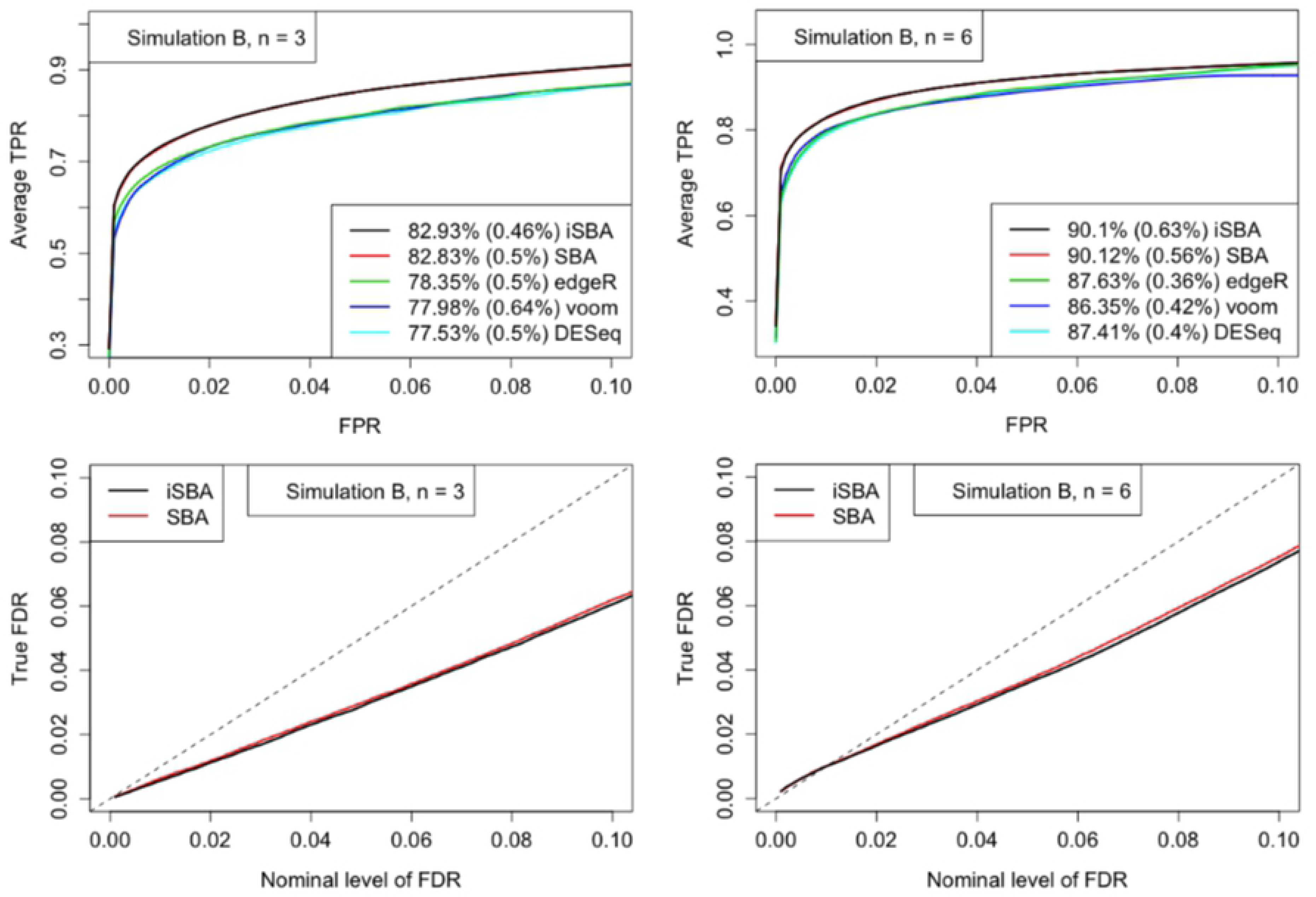
Results for testing *|logFC| ≤ log*1.5 from Simulation B. The upper panel shows the ROC curves. For each level of FPR, the TPRs were averaged over the 32 simulated datasets. The percentage reported in the legend is the average AUC for each method, representing the percentage of 0.1, which is the total area in the range of FPR *<* 0.1, and the percentage in each set of parentheses is the standard deviation of the estimated AUC. The lower panel plots the actual FDR versus the nominal level of FDR. The dashed lines correspond to the *Y* = *X* line.

### Simulation Results for Swapping Treatments

As we discussed in the Introduction Section, the semi-parametric Bayesian (*SBA*) method [13] set one treatment group as reference condition. If the choice of a reference condition is not obvious based on the experimental design, the declared differential expression status may vary depending on which group is set to be baseline. However, the model we proposed is invariant no matter which group is set to be the reference condition. The proportion of genes remaining the same declared differential expression status between two analyses that swapped the treatment and control groups were calculated when controlling FDR at 0.05, with average values and standard deviation of the percentage across the 32 simulated datasets reported in Table 1. It turned out that our *iSBA* method had higher overlap and more consistency in declared differential expression status than *SBA* method when swapping the treatment and control groups, for all simulation settings.

**Table 1.** Proportion of genes remaining the same declared differential expression status between two analyses that swapped the treatment and control groups for Simulations A and B, when controlling FDR at 0.05. The proportions were averaged across the 32 simulated datasets, and the percentage in each set of parentheses is the standard deviation of the estimated proportion.

| Simulation setting | *SBA* | *iSBA* |
| --- | --- | --- |
| Simulation A, $n = 3$ | 91.43% (2.21%) | 92.43% (0.33%) |
| Simulation A, $n = 6$ | 93.12% (5.55%) | 94.87% (0.48%) |
| Simulation B, $n = 3$ | 91.42% (4.80%) | 93.78% (0.84%) |
| Simulation B, $n = 6$ | 93.98% (2.23%) | 94.83% (0.39%) |

### Real Data Analysis

In this subsection, we analyze a real RNA-seq dataset published by Li et al. (2010). The dataset measures the transcript abundance of two cell types, bundle sheath and mesophyll, for different leaf sections. Each cell type has two biological replicates. The objective of the analysis is to detect genes that are DE between cell types or between different leaf sections. We analyzed leaf section 4 to detect DE genes between the two cell types in this section.

After deleting genes that have zero counts for both replicates in either cell type, 28,407 out of 33,743 genes were retained for analysis. We assumed NB models for the count data observed for each gene, and performed our proposed *iSBA* method, together with *SBA* method and *edgeR*. We also controlled FDR as described in subsection “Bayesian FDR Control” for *SBA* and *iSBA*, and applied the Benjamini and Hochberg [23] procedure for *edgeR*.

The numbers of DE genes detected by different methods while controlling FDR at different levels are shown in Fig 5. For example, when we controlled FDR at 0.05, 6040 genes were detected by all three methods. The majority of genes identified by our *iSBA* method were overlapped with *SBA*. 2703 genes were detected by both *iSBA* and *SBA*, but not by *edgeR*, which may due to the conservative control of FDR based on our simulation studies.

**Fig 5.**
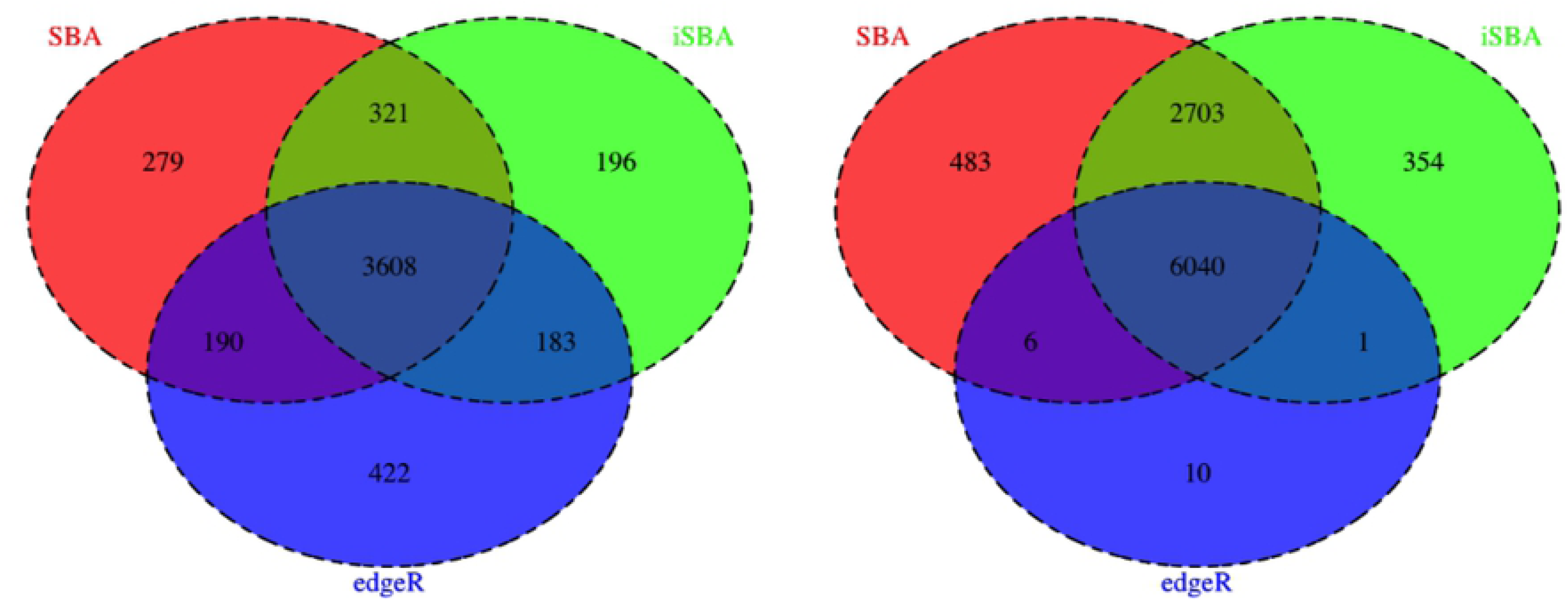
The numbers of DE genes between two cell types for leaf section 4. The Venn diagram on the left shows the number of overlapping identified DE genes from our *iSBA* method, *SBA* method, and *edgeR* while controlling FDR at 1%; the Venn diagram on the right shows the corresponding results while controlling FDR at 5%.

The proportions of genes remaining the same declared differential expression status between two analyses that swapped the two treatment groups when controlling FDR at for our *iSBA* method is 93.47%, while the *SBA* method is 89.28%.

## Discussion

In this paper, we proposed a Bayesian mixture modeling procedure for DE analysis of RNA-seq count data, and employed the MCMC sampling scheme to generate posterior samples for further inference. Simulation results demonstrate that our method outperformed other commonly used methods, such as *edgeR, voom* and *limma* pipeline, and *DESeq*, in terms of both ranking DE genes and FDR control.

A common choice of the concentration parameter *M* in the DP priors that are widely used in application is *M* = 1 [12]. We check the simulation results with different *M* values (*M* being 0.2, 0.5, 2, 5, 10 or 20), and the results remain almost the same for various values of *M*.

In our proposed method, the DP prior we choose guarantees that our modeling is invariant regardless of which treatment group is set to be the reference condition. According to the simulation results on two analyses that swapped the treatment and control groups, it is worth noticing that even for our *iSBA* method, the declared differential expression status are still not 100% the same. Part of the reason is due to the randomness of MCMC, if we run another MCMC using different seed, the overlap between the two MCMCs is about 97%. Since we generated 5000 posterior samples to calculate the estimated posterior probabilities after 3000 iterations burn-in, whether the Markov chains are long enough to get accurate results is also a potential problem. We checked the effective sample size for each gene, genes that had the same declared DE status after swapping treatments had effective sample sizes about 500 or larger, but genes that had different declared DE status overlapped had effective sample sizes around only 100. Effective sample size around 400 can be regarded as large enough, so for those genes with low effective sample size, we may need to run longer MCMC. Based on simulation checking, running the Markov chains longer do increase the percentage of overlapping genes, as expected. For example, for simulation A with *n* = 6, if we doubled the length of chain, the overlap for *iSBA* increased to 95.28%. However, running longer chains is more time consuming, and it only benefits a small proportion of genes while results for most genes would not change. Therefore, it is a tradeoff between efficiency and accuracy, and we will let the users decide which one is more important for a practical application.

As indicated in subsection “Bayesian FDR Control”, the estimated posterior probability 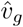 is used as a test statistic and a decision *D*_*g*_ is based on whether 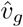 is small enough. And the AMAP test by Si and Liu (2013) [11] is based on a similar test statistic except that the prior models are different. In fact, the MAP test statistic derived in Si and Liu (2013) [11] can be viewed as the posterior probability of being null given the distribution of gene-specific parameters under the null hypothesis and the distribution of these parameters under the alternative hypothesis. Assuming the distributions of the gene-specific parameters (fold changes and other parameters) follow approximately the prior distribution we use, our estimated posterior probability using MCMC is an AMAP test statistic.

## Acknowledgments

The authors would like to thank Emily Goren from Iowa State University for proofreading the manuscript.

## Supporting information

**S1 Appendix. Proof of Model Invariance.**

**S2 Appendix. Detailed MCMC Sampling Scheme.**

